# Mechanical Stiffening Promotes Growth, Invasion, and Mitogen-Activated Protein Kinase Kinase (MEK) Inhibitor Resistance in 3D Plexiform Neurofibroma Cultures

**DOI:** 10.64898/2026.01.26.701794

**Authors:** Kyungmin Ji, Chenjun Shi, Jitao Zhang

## Abstract

Plexiform neurofibromas with neurofibromatosis type I (pNF1s) are benign peripheral nerve sheath tumors caused by *NF1* loss, leading to dysregulated RAS/mitogen-activated protein kinase (MAPK) signaling. While mitogen-activated protein kinase kinase (MEK) inhibitors, selumetinib and mirdametinib, can reduce tumor volume, surgical resection remains the primary treatment for immediate debulking and symptom relief. Complete removal is often limited by tumor infiltration along nerve plexuses, and residual tumors may undergo postsurgical tissue remodeling, producing localized regions of stiffened extracellular matrix (ECM). The impact of ECM stiffness on pNF1 progression and drug response is unknown. Using patient-derived pNF1 tumor cells cultured in 3D hydrogels with defined stiffness (1.5 kPa soft, 7 kPa stiff), we found that stiff ECM promoted spread morphology, increased growth, and progressive intracellular softening. Stiff ECM reduced lysyl oxidase (LOX) expression, reflecting mechanoadaptive ECM remodeling, and upregulated P-glycoprotein, leading to decreased sensitivity to selumetinib. These results provide the first evidence that ECM stiffening, such as that arising from postsurgical remodeling, directly drives pNF1 progression and therapeutic resistance. Our findings highlight mechanobiology as a key regulator of tumor behavior and support targeting ECM mechanics to improve clinical outcomes in NF1 patients.

## Introduction

Plexiform neurofibromas (pNF1s) are benign, diffuse peripheral nerve sheath tumors that arise in individuals with neurofibromatosis type I and are driven by loss of neurofibromin, a key negative regulator of RAS/mitogen-activated protein kinase (MAPK) signaling[1-6]. These tumors grow diffusely along nerve plexuses and infiltrate surrounding soft tissues, often reaching substantial size and causing pain, functional impairment, and disfigurement[1-6]. Complete surgical removal is frequently difficult because of their infiltrative growth along nerves, blood vessels, and other critical organs, leading to residual tumor tissue and the need for repeated surgical interventions[1-6]. Although histologically benign, approximately 8–13% of pNF1s progress to malignant peripheral nerve sheath tumors (MPNSTs), which are aggressive and carry poor prognoses[7]. FDA-approved MEK inhibitors, selumetinib (AZD6244) [8,9] and mirdametinib (PD0325901) [10-12], can reduce tumor volume by ∼30% and are used for inoperable, progressive, or symptomatic tumors, particularly when resection is incomplete or associated with high morbidity[13,14]. However, therapeutic responses are partial, and residual tumor regions may form mechanically distinct microenvironments that influence cellular signaling and drug efficacy. As a result, surgical resection remains [15,16]the primary treatment when feasible, providing immediate tumor debulking and symptom relief. However, because of their infiltrative growth and involvement with nerves, blood vessels, and other vital structures, pNF1s often persist or recur after incomplete surgical resection, necessitating repeated surgeries over a patient’s lifetime[17-19].

ECM stiffening is increasingly recognized as a critical regulator of tumor biology in other solid cancers [20-22]. Increased matrix rigidity can arise from enhanced deposition and crosslinking of collagen and other ECM components, often mediated by enzymes such as lysyl oxidase (LOX), and is frequently associated with tumor progression, fibrosis, or chronic tissue remodeling [22-24]. ECM stiffening can also occur as part of wound healing after surgery, when fibroblast activation and collagen deposition locally increase matrix rigidity[25-29]. Stiffened ECM enhances cell proliferation, promotes invasive behavior, and reduces drug efficacy through mechanotransduction pathways involving integrins, focal adhesion kinase (FAK), and yes-associated protein (YAP)/ transcriptional co-activator with PDZ-binding motif (TAZ) signaling [20,23,30]. In addition, ECM stiffening influences cytoskeletal organization and intracellular mechanics, creating a mechanically distinct microenvironment that feeds back to regulate tumor cell behavior [30-32]. Because residual pNF1 tumors persist after surgery, it is plausible that postsurgical tissue remodeling—including localized ECM stiffening—occurs, although this has not been systematically characterized. Understanding how ECM stiffness influences pNF1 growth, invasion, and drug responsiveness could uncover mechanisms driving tumor progression and therapy resistance, providing clinically relevant insights to guide treatment strategies for NF1 patients.

To address this knowledge gap, we developed a three-dimensional (3D)/4D (3D in real-time) hydrogel-based culture system in which patient-derived pNF1 tumor cells are embedded in matrices of precisely tuned stiffness. This system models ECM conditions ranging[22,33] from compliant tissue to fibrotic, stiffened regions, allowing controlled assessment of stiffness-dependent effects on tumor growth, ECM remodeling, and drug response. Understanding ECM stiffness is particularly important in pNF1, as surgical resection - the mainstay treatment[15,16] - often leaves residual tumors, and postsurgical remodeling can locally stiffen the matrix. Our platform thus enables investigation of how mechanically altered microenvironments in residual tumors may drive tumor progression, infiltrative behavior, and reduced therapeutic sensitivity, providing a mechanistic framework for mechano-aware strategies in NF1.

## Materials and Methods

### Reagents and antibodies

LunaX™ PureMatrix – photocrosslinkable ECM with low or high stiffness and LunaX™ crosslinker were purchased from Gelomics (Kelvin Grove, Queensland, Australia). Phenol red-free Dulbecco’s Modified Eagle Medium (DMEM) and MycoZap™ Plus-CL were purchased from Lonza (Basel, Switzerland). Fetal bovine serum (FBS) was obtained from Cytiva (Marlborough, MA). Selumetinib (AZD6244) was purchased from Selleckchem (Houston, TX, USA). L-glutamine and all other chemicals, unless otherwise specified, were purchased from Sigma-Aldrich (St. Louis, MO). LIVE/DEAD™ Viability/Cytotoxicity kit, actin-phalloidin, Hoechst33342 and 4′-6-Diamidino-2-phenylindole (DAPI) were purchased from ThermoFisher Scientific (Waltham, MA). Antibodies against lysyl oxidase (LOX) and p-glycoprotein (Pgp) were purchased from Abcam (Cambridge, UK).

### Cell lines and cell maintenance

Two immortalized human plexiform neurofibroma cell lines (*Nf1*^*-/-*^; hereafter referred to as pNF1 tumor cells), ipNF95.11bC and ipNF05.5, were obtained from ATCC and previously described [34-36]. Cells were maintained as monolayers in DMEM/high glucose supplemented with 10% FBS at 37°C, 5% CO_2_. Routine RT-PCR screening confirmed the absence of mycoplasma contamination.

### Three-dimensional (3D)/4D (3D in real-time) culture models

3D cultures were established in our patented microfluidic culture devices called TAME (tissue architecture and microenvironment engineering; Patent#, US10227556B2) chips[37] as previously published[35,36,38]. Fabrication of TAME chips, including separate and linked well designs[37], and protocols for 3D cultures have been described previously [35,36,39]. Embedding of cells in LunaX™ PureMatrix – photocross-linkable ECM with low or high stiffness and the crosslinking of 3D hydrogel with different stiffness were followed by the manufacturer’s instructions[40,41]. Briefly, 50,000 cells were mixed with 50 ml of 3D hydrogel with low or high stiffness and 50 ml of photoinitiator, plated in the glass-bottom wells of TAME chips and exposed to LunaX™ crosslinker for 50 sec. Then cells with DMEM supplemented with 5% FBS were maintained for the indicated culture periods (3-10 days). Prior to live-cell 3D imaging, nuclei were stained with Hoechst 33342 to assess cell integrity and morphology, and total live-cell numbers were quantified using viability markers such as Calcein-AM.

### Brillouin microscopy

The mechanical properties of cells were quantified by using a confocal Brillouin microscope. The detailed configuration of this microscope can be found from previous publication[42]. Briefly, a 660-nm continuous-wave laser was used as light source. The laser beam was coupled into a commercial optical microscope (IX83, Olympus) and focused into the sample using an objective lens (0.6 NA, Olympus), yielding a beam spot of 0.7 × 0.7 × 2.5 µm. The back-scattered Brillouin signal was collected by the same objectives and sent to a Brillouin spectrometer for analysis. During experiments, 10-20 mW laser power was shined onto a cell, and Brillouin image of the cell was collected with an acquisition speed of 50 ms per pixel and a step size of 1 µm. The Brillouin shifts of all pixels was then averaged to represent the stiffness of each cell.

The confocal Brillouin microscopy is based on the optical process known as spontaneous Brillouin scattering, in which the scattered light experiences an optical frequency shift, called Brillouin shift, due to the interaction of the incident light and the material. In backward-scattering configuration, the elastic longitudinal modulus *M*′ is related to the Brillouin shift *ω*_*B*_ by

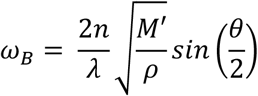

where *λ* is the wavelength of incident light, *n* and *ρ* are refractive index and density of the material. For some biological materials such as cells, the ratio of 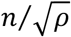 can be approximated as constant [43]. Therefore, the measured Brillouin shift can be used to estimate the relative change of mechanical modules.

### Live-cell LIVE/DEAD™ Viability/Cytotoxicity assay

This assay was performed as described in our previous studies[36,38], and is summarized briefly here. 3D cultures were incubated with Calcein AM (live cells, green) for 30 minutes at 37 °C, followed by a single wash with warm PBS. Fresh warm culture media were then added according to the manufacturer’s instructions and previously published protocols [35,36,38]. Prior to live-cell imaging, nuclei were stained with Hoechst 33342 to assess nuclear morphology and confirm cell viability. Live-cell imaging was performed using confocal microscopy under physiological conditions. Quantification of viable cells was conducted in 3D using Volocity software by counting nuclei that were intact, spherical, and evenly stained (excluding condensed or fragmented nuclei) and colocalized with Calcein AM-positive cytoplasm.

### Image acquisition for quantitative analysis in 3D

As described in our prior studies[35-37,44], live-cell imaging and quantitative analysis were performed in 3D. Optical sections spanning the entire depth of the 3D cultures were acquired from four contiguous fields (2 × 2 or 3 × 3) using either an upright Zeiss LSM 780 confocal microscope (Carl Zeiss, Jena, Germany) equipped with a temperature- and CO_2_-controlled environmental chamber, or an inverted Leica Stellaris 5 system (Leica Microsystems, Deerfield, IL, USA) with a Tokai Hit stage-top incubator to maintain physiological conditions. Image stacks were processed and reconstructed in 3D using Volocity software (Version 7.0.0; PerkinElmer, Waltham, MA, USA) to visualize and quantify viable cell distributions. In reconstructed volumes, spatial orientation is indicated by arrows showing the x-axis (green), y-axis (red), and z-axis (blue) in the lower corner of each image.

### Immunofluorescence staining

pNF1 tumor cells were cultured for 5 days in 3D hydrogels of either soft or stiff ECM. Following culture, cells were fixed with 10% formaldehyde for 10 minutes at room temperature (RT), washed with PBS, and permeabilized for 5 minutes using 0.2% Triton X-100 in PBS. Cells were then blocked with 1% BSA for 1 hour at RT and incubated with primary antibodies against LOX or P-glycoprotein for 3 hours at RT. After washing, cells were incubated with appropriate fluorescent secondary antibodies for 1 hour at RT. Nuclei were counterstained with DAPI. Images were acquired using an inverted Leica Stellaris 5 confocal microscope (Leica Microsystems, Deerfield, IL, USA). Quantification of LOX and P-glycoprotein fluorescence intensity was performed using ImageJ software (National Institutes of Health, Bethesda, MD, USA).

### Statistical analysis

Data are presented as bar graphs or box-and-whisker plots. In bar graphs, bars represent the mean ± standard deviation (SD). In box-and-whisker plots, boxes represent interquartile ranges and whiskers minimum and maximum values. The number of replicates is provided in each corresponding figure legend. Statistical significance was assessed using either one-way ANOVA followed by Tukey’s post-hoc test for multiple group comparisons, or Student’s t-test for comparisons between two groups. A *p*-value ≤ 0.05 was considered statistically significant throughout the study.

## Results

### Stiff ECM promotes spread morphology and enhances 3D growth of pNF1 tumor cells

Because pNF1s often persist or recur after incomplete surgical resection [17-19] and postsurgical remodeling may stiffen the ECM[25-29], we first examined how matrix stiffness affects tumor cell behavior. Two patient-derived immortalized pNF1 cell lines, ipNF95.11bC and ipNF05.5, were cultured in 3D hydrogels with defined stiffness values of 1.5 kPa (soft) and 7 kPa (stiff). These stiffness values were selected based on prior mechanobiology studies in multiple tumor models [22,33], where matrices in the ∼1–2 kPa range are commonly used to represent physiologically soft tissues, whereas matrices in the ∼5– 10 kPa range are widely employed to model fibrotic or mechanically stiffened tumor microenvironments, allowing us to model both compliant and fibrotic tumor conditions.

After 10 days, cells were stained with phalloidin to visualize filamentous actin and counterstained with Hoechst 33342 for nuclei. 3D reconstruction in Volocity revealed that cells in soft ECM maintained a compact and rounded morphology, whereas cells in stiff ECM adopted a more spread and expanded shape, reflected by a significant increase in the longest axis length in 3D (Fig. 1A–D). Volumetric reconstruction further demonstrated that total cell number was significantly higher in stiff ECM compared with soft ECM for both cell lines (p < 0.01) (Fig. 1E–F), indicating that ECM stiffness alone promotes pNF1 tumor growth and morphological expansion.

**Fig. 1.**
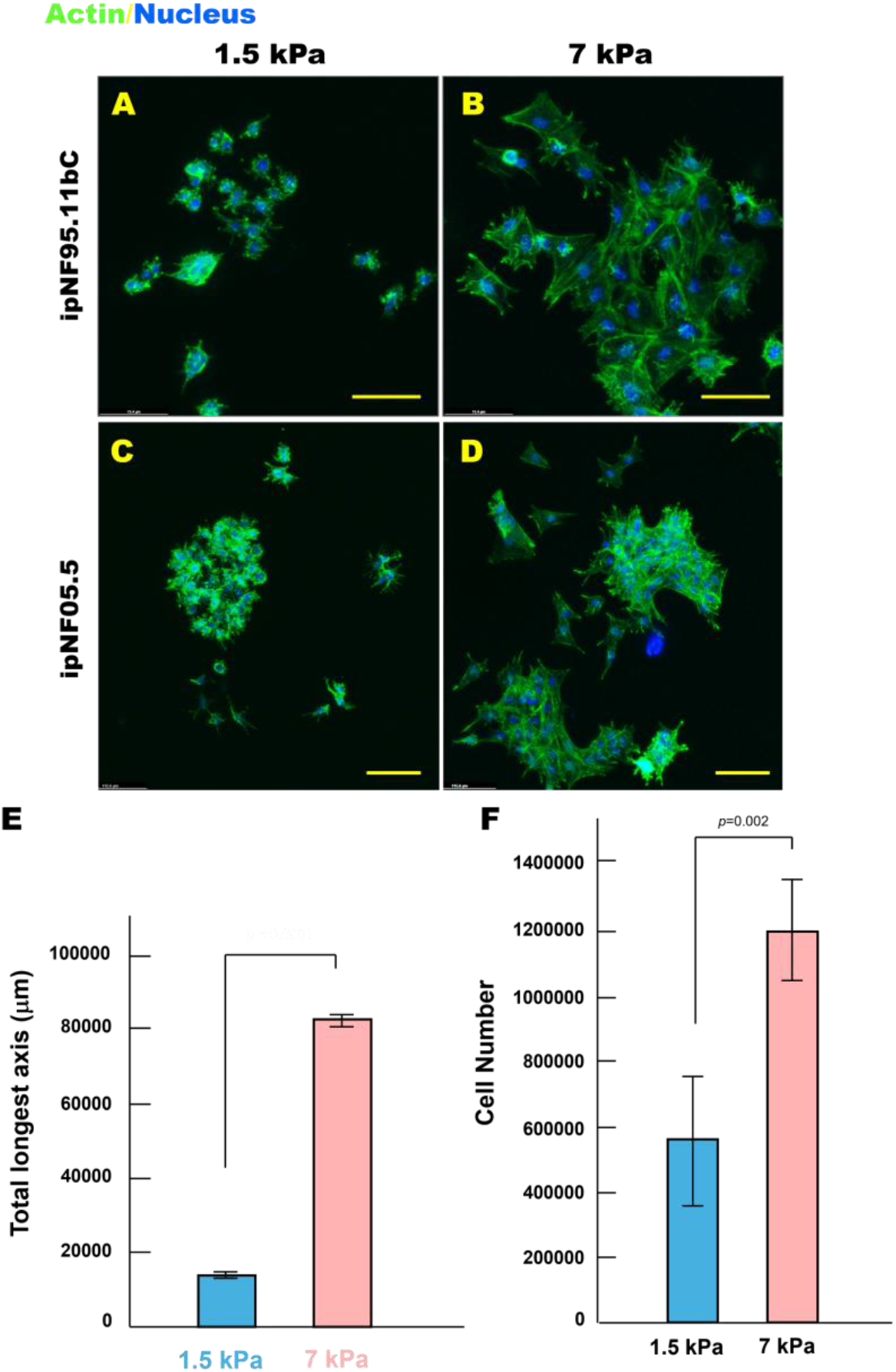
pNF1 tumor structures exhibit a more spread morphology and increased growth in stiff ECM compared with soft ECM in 3D culture. (A-D) En face views of 3D reconstructions of pNF1 structures formed by ipNF95.11bC (A, B) and ipNF05.5 (C, D) monocultures grown in soft (1.5 kPa) or stiff (7 kPa) ECM for 10 days. Actin fibers (green, phalloidin) and nuclei (blue) are shown. Scale bars: 70 µm (top row) and 115 µm (bottom row). (E, F) Quantification of structure size (longest axis, E) and cell number (F) measured in 3D using Volocity. Data represent mean ± SD (n = 4).

### Stiff ECM Induces progressive intracellular softening in 3D pNF1 tumor structures

Because decreased cellular stiffness has been linked to enhanced deformability and infiltrative behavior in multiple solid tumors[45-48], we next measured intracellular stiffness in our 3D pNF1 cultures to assess whether ECM mechanical cues similarly affect infiltrative potential. Intracellular stiffness was quantified using Boullion microscopy[49,50], a non-invasive, high-resolution technique that enables real-time, quantitative measurement of cytosolic mechanics in living 3D cultures without disrupting tissue architecture. Cells were cultured in soft (1.5 kPa) or stiff (7 kPa) ECM for 9 days. Cells in soft ECM maintained relatively stable intracellular stiffness throughout the culture period (the average Brillouin shifts were within the range of 6.44 GHz and 6.48 GHz) whereas cells in stiff ECM exhibited progressive cytosolic softening over time (the average Brillouin shift dropped from 6.50 GHz on Day 7 to 6.32 GHz on Day 9) (Fig. 2A–C). This stiffness-dependent intracellular adaptation suggests that mechanical cues from the ECM induce increased cellular deformability, potentially facilitating enhanced infiltration into surrounding matrix regions. These results set the stage for examining whether intracellular softening is coupled to extracellular matrix remodeling.

**Fig. 2.**
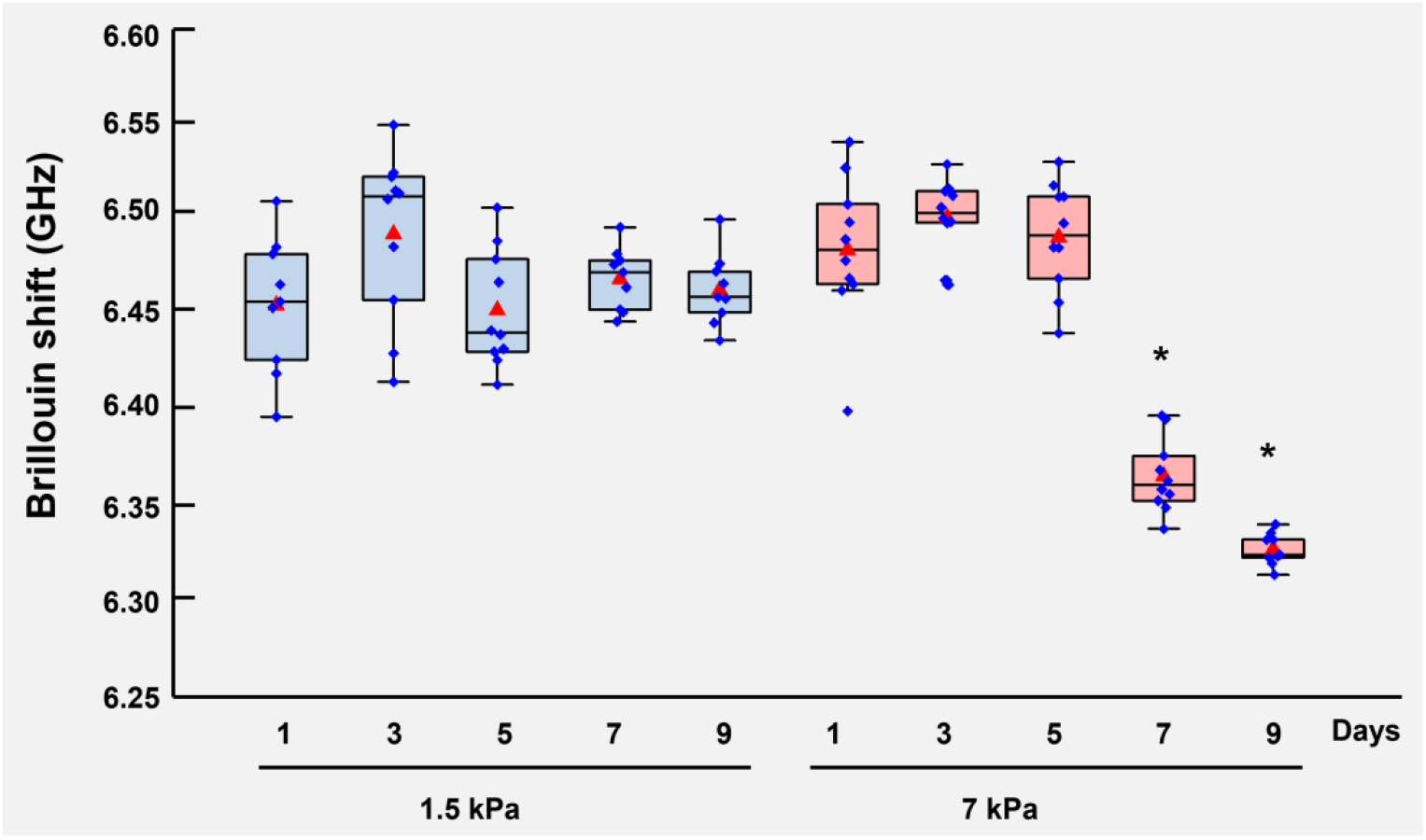
Stiff ECM induces progressive intracellular softening in 3D pNF1 tumor structures. pNF1 tumor cells, ipNF95.11bC, were embedded in soft (1.5 kPa) or stiff (7 kPa) 3D ECM hydrogels and cultured for 9 days. Intracellular cytosolic stiffness was quantified at the indicated times using Brillouin microscopy. Individual cell bodies were scanned longitudinally over time to assess stiffness adaptations in response to ECM mechanics. * *p* < 0.01 *vs*. Day 1 (n=3).

### ECM stiffness regulates LOX-mediated matrix remodeling

Because intracellular mechanical adaptation may influence interactions with the surrounding matrix, we measured lysyl oxidase (LOX) expression, an enzyme that crosslinks collagen and elastin to regulate ECM stiffness[22-24]. pNF1 tumor cells were cultured for 10 days in soft or stiff ECM, and LOX expression was quantified by immunofluorescence. Tumor structures in soft ECM expressed significantly higher levels of LOX than those in stiff ECM (Fig. 3A–C). This suggests a mechanoadaptive feedback mechanism[51] in which pNF1 tumor cells upregulate LOX in compliant matrices to reinforce ECM rigidity, whereas stiff matrices do not require additional LOX-mediated crosslinking. Importantly, these findings link directly to our intracellular softening results: stiff ECM induces cytosolic softening while simultaneously reducing LOX expression, highlighting coordinated mechanical adaptation at both cellular and extracellular levels.

**Fig. 3.**
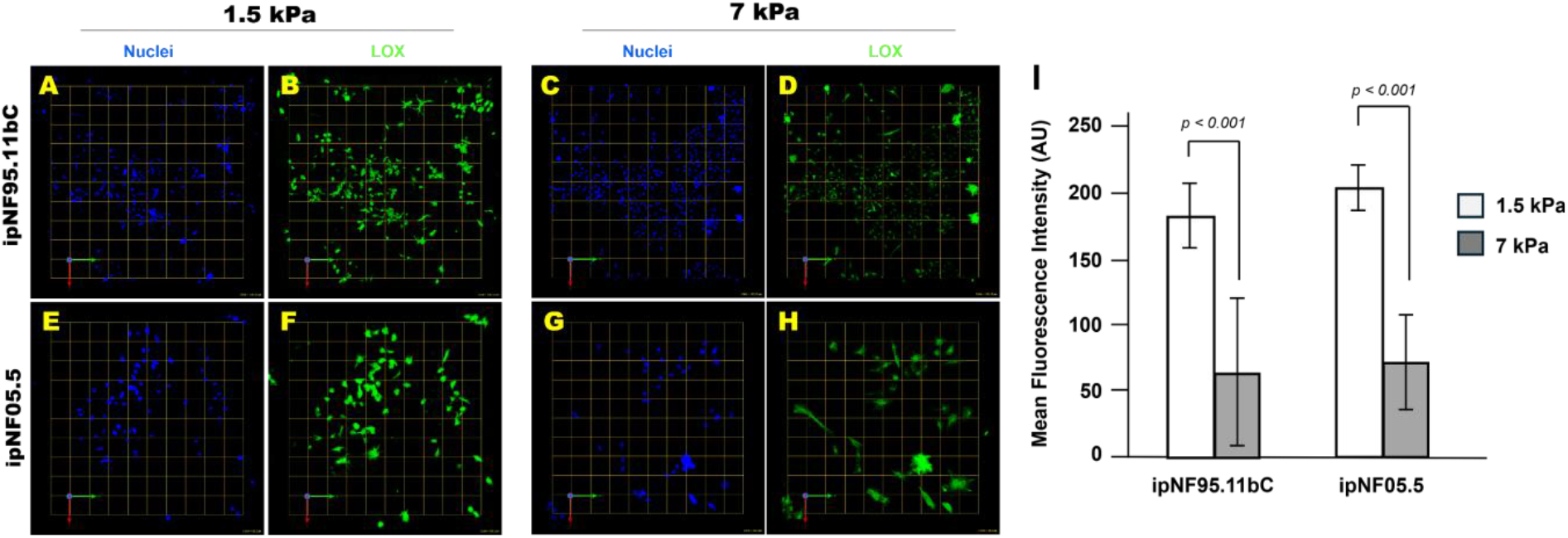
Stiff ECM downregulates LOX expression in 3D pNF1 tumor structures. (A-H) *En face* views of 3D reconstructions of pNF1 tumor structures formed by ipNF95.11bC (A-D) and ipNF05.5 (E-H) cultured in soft (1.5 kPa) or stiff (7 kPa) 3D ECM for 10 days. LOX expression is shown in green, and nuclei are labeled in blue. Each grid square corresponds to 110 µm (A-D) or 58 µm (E-H). (I) Quantification of LOX fluorescence intensity was performed using ImageJ. Data represent mean ± SD (n = 3).

### Stiff ECM enhances p-glycoprotein expression and reduces sensitivity to selumetinib

Finally, to determine whether stiffness-driven intracellular and extracellular adaptations have functional consequences for therapy, we evaluated the effect of ECM stiffness on drug resistance. Mechanotransduction and ECM mechanics regulate the expression and activity of drug-efflux transporters [e.g., ATP-binding cassette subfamily B member (ABCB) 1/p-glycoprotein (Pgp), ATP-binding cassette subfamily C member (ABCC) 1/ multidrug resistance-associated protein (MRP) 1 and ATP-binding cassette subfamily G member (ABCG) 2/ breast cancer resistance protein (BCRP)] via integrin–FAK/ILK and YAP/TAZ-dependent pathways, and this link has been demonstrated in multiple tumor models including breast, lung, colorectal and others [52-55], highlighting its clinical relevance for understanding partial responses to selumetinib in pNF1 patients. Tumor cells were cultured for 10 days in soft or stiff ECM and analyzed for Pgp expression by immunofluorescence. Tumor structures in stiff ECM exhibited significantly higher Pgp levels than those in soft ECM (Fig. 4). To assess whether this stiffness-associated upregulation of drug efflux affects therapeutic response, tumor cells were treated with selumetinib for 5 days. Live and dead cells were labeled with Calcein-AM and ethidium homodimer-1, respectively, and total cell viability was quantified using Volocity. Tumor structures in stiff ECM retained substantially more viable cells after treatment compared with those in soft ECM, indicating reduced sensitivity to MEK inhibition (Fig. 5).

**Fig. 4.**
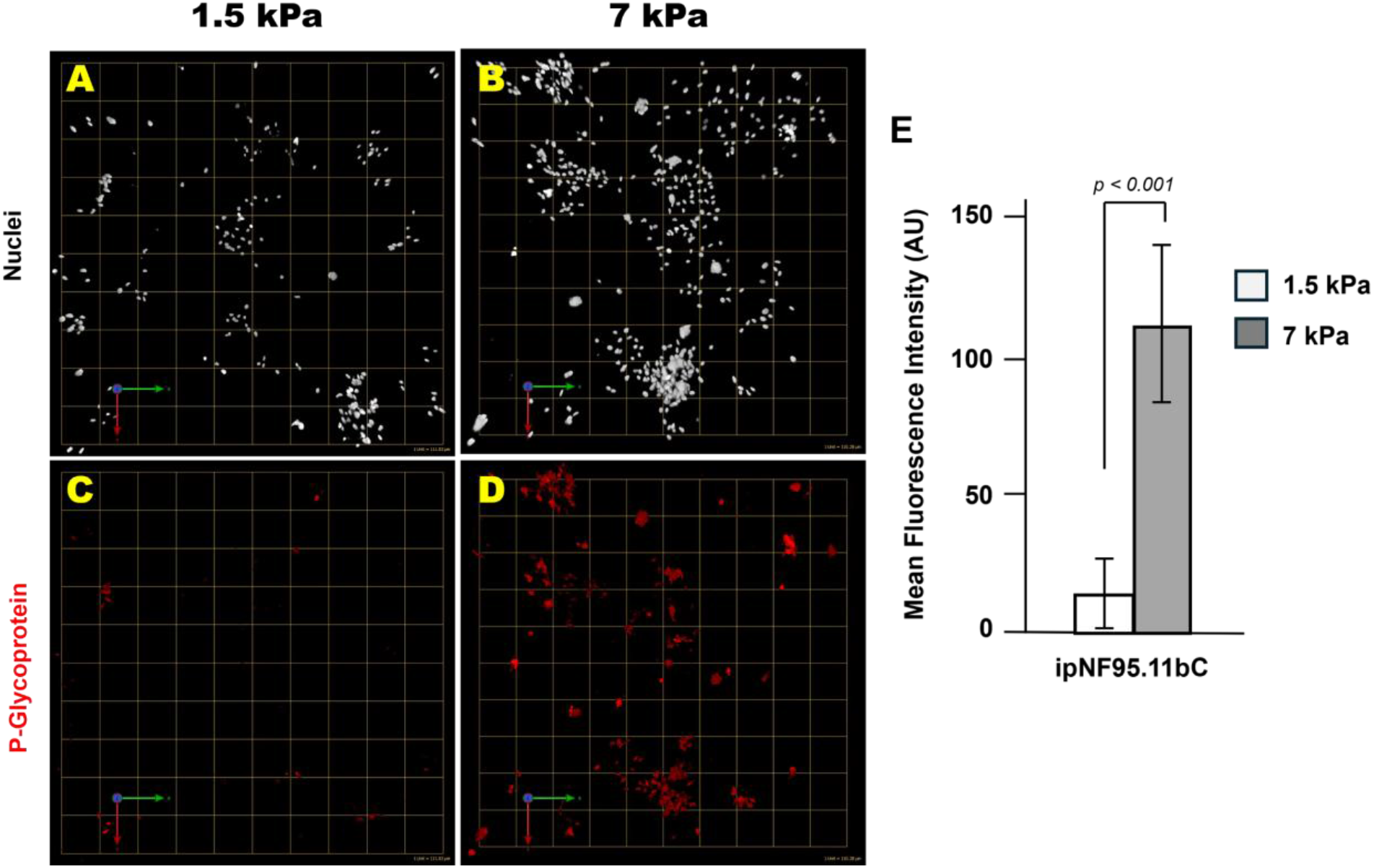
Stiff ECM promotes Pgp expression in 3D pNF1 tumor structures. (A–D) *En face* views of 3D reconstructions of pNF1 tumor structures formed by ipNF95.11bC cultured in soft (1.5 kPa) or stiff (7 kPa) 3D ECM for 10 days. Pgp expression is shown in red, and nuclei are labeled in pseudowhite. Each grid square corresponds to 110 µm. (E) Quantification of Pgp fluorescence intensity was performed using ImageJ. Data represent mean ± SD (n = 3).

**Fig. 5.**
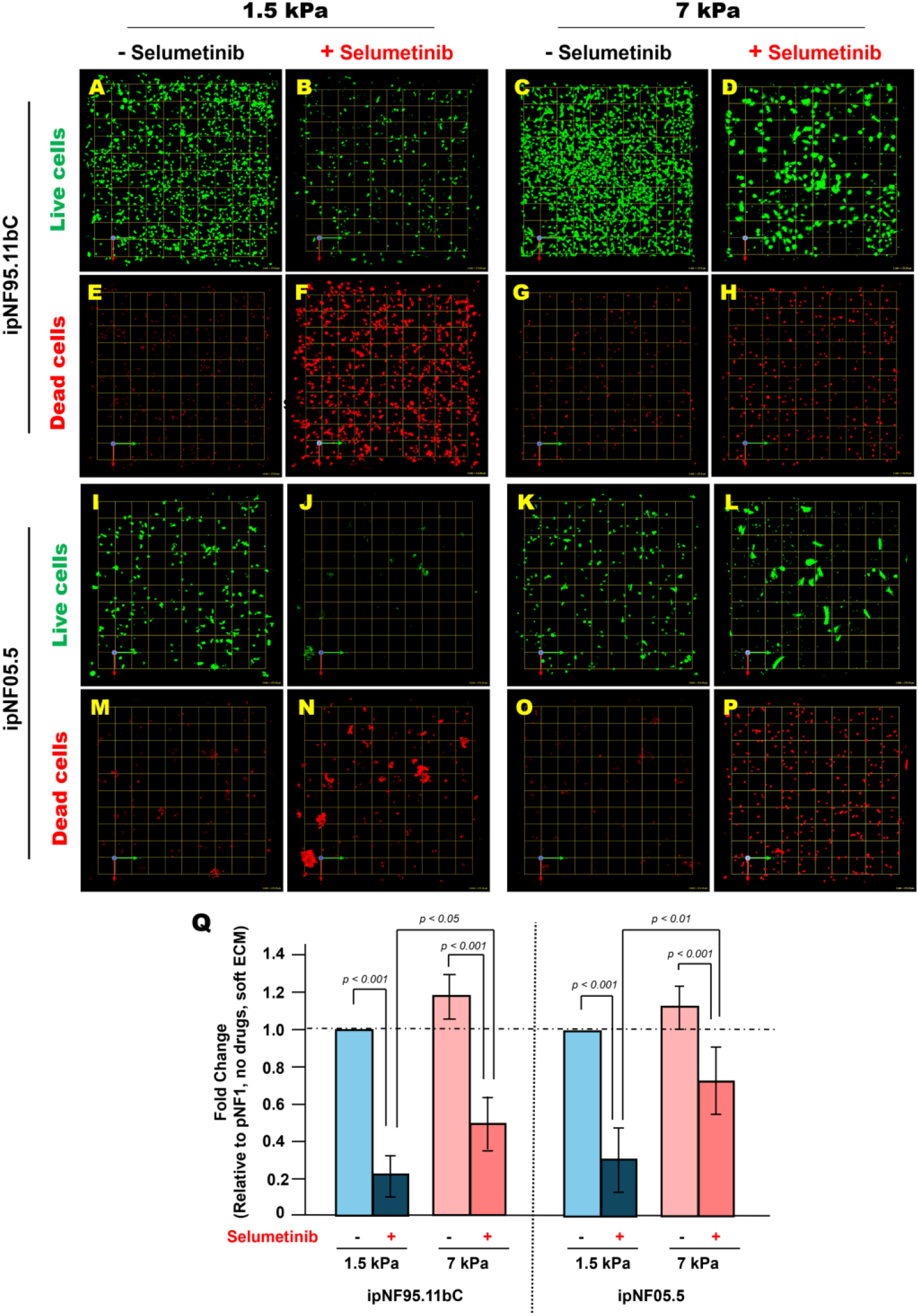
Stiff ECM increases drug resistance in 3D pNF1 tumor structures after 5-day selumetinib treatment. (A-P) *En face* views of 3D reconstructions of pNF1 tumor structures formed by ipNF95.11bC(A-H) and ipNF05.5 (I-P) in monocultures without (first and third panels) or with 5-day selumetinib treatment (second and fourth panels). Live and dead tumor cells were stained with Calcein-AM (green) and ethidium homodimer-1 (red), respectively. Images are tiled from 4 contiguous fields. Grid, 174 µm. (Q) Quantification of total live pNF1 tumor cells in 3D monocultures using Volocity. Data are expressed as fold change relative to untreated cells in soft ECM. Each bar represents mean ± SD (n = 3).

Collectively, these results demonstrate that stiff ECM enhances multiple pNF1 phenotypes, including growth, spread morphology, intracellular softening, reduced LOX expression, and decreased drug sensitivity. This integrated response highlights the critical role of mechanical cues in modulating tumor behavior and therapy resistance in pNF1s.

## Discussion

This study demonstrates that ECM stiffness is a critical, previously underappreciated determinant of pNF1 tumor cell behavior. Using a tunable 3D hydrogel system combined with quantitative mechanophenotyping, we show that mechanical rigidity alone - independent of biochemical cues - is sufficient to regulate multiple dimensions of pNF1 tumor biology, including morphology, intracellular mechanics, ECM remodeling, and response to targeted therapy. These findings expand the framework of pNF1 biology beyond canonical RAS/MAPK hyperactivation and identify ECM stiffness as a potential driver of residual tumor progression and therapy resistance.

A major and novel observation is that stiff ECM promotes spread morphology and enhances growth of pNF1 tumor structures. Clinically, this is particularly relevant because surgical resection remains the primary treatment for pNF1[15,16], yet complete removal is limited by the diffuse and infiltrative nature of these tumors[17-19]. Residual tumor tissue may reside within mechanically rigid, fibrotic niches[25-29], which could favor regrowth and expansion, providing a mechanistic explanation for postoperative recurrence.

At the cellular level, stiff or stiffening ECM environments can elicit mechanoadaptive responses in which tumor cells become more deformable or exhibit reduced intracellular stiffness [56,57]. Our findings with pNF1 tumor cells align with this behavior. Such decreased stiffness has been consistently associated with enhanced deformability and greater infiltrative potential across multiple solid tumor models [45-48,58]. Although pNF1s are histologically benign, they grow diffusely along nerve pathways [1-6], and our data suggest that stiffness-induced cytoskeletal softening may help enable this infiltrative behavior, revealing a previously unrecognized mechanoadaptive response in pNF1s. ECM rigidity also altered expression of lysyl oxidase (LOX), a key collagen- and elastin-crosslinking enzyme [22-24]. We observed higher LOX expression in soft ECM and reduced LOX in stiff ECM, a pattern consistent with mechano-responsive feedback loops described in other tissues and tumor systems [51]. These findings indicate coordinated intracellular and extracellular adaptation, in which pNF1 tumor cells remodel both their cytoskeleton and matrix-regulatory programs in response to ECM stiffness.

Our prior studies[35,36] demonstrated that fibroblast-derived secretome enhances pNF1 tumor cell growth, infiltrative behavior, and drug resistance in 3D cultures. In patient tumors, tumor cells comprise only ∼6% of total volume, whereas fibroblasts represent ∼60% and are the primary collagen producers[59]. Moreover, fibroblasts are more resistant to MEK inhibitors than tumor cells in 3D[36]. Based on these observations, it is plausible that fibroblasts in residual tumors—after surgery or drug treatment—could contribute to localized ECM stiffening. Such mechanically rigid niches may promote tumor expansion, enhanced infiltration, and reduced therapeutic sensitivity, providing a mechanistic link between stromal remodeling and pNF1 progression.

Stiff ECM also increased P-glycoprotein (Pgp) expression and reduced sensitivity to MEK inhibitors, including selumetinib and mirdametinib, indicating that mechanical cues in the tumor microenvironment can directly influence drug response. Notably, our prior studies[36] demonstrated that fibroblast-derived secretome upregulates Pgp expression and enhances drug resistance in pNF1 tumor cells. These observations suggest that both stromal signals from fibroblasts and mechanically stiffened ECM may converge on Pgp-mediated mechanisms to modulate therapeutic sensitivity. Given that surgical resection remains the primary treatment for pNF1s [15,16] and clinical responses to MEK inhibition are often partial [13,14], these findings underscore the translational relevance of considering ECM mechanics and stromal contributions in evaluating residual tumor behavior and optimizing therapy.

The 3D hydrogel-based model represents a major methodological advance for pNF1 research. By enabling precise tuning of ECM stiffness while preserving native 3D architecture, this platform allows controlled assessment of stiffness-dependent growth, cytoskeletal mechanics, ECM remodeling, and drug response. Building on our prior 3D coculture work incorporating bioengineered neuronal axons[38], this system demonstrates versatility and can be further adapted to study additional microenvironmental interactions, such as tumor–axon crosstalk, alongside mechanobiology. Importantly, it provides a physiologically relevant framework for preclinical testing of mechanotransduction-targeted therapies, LOX inhibitors, or antifibrotic approaches, and may facilitate incorporation of mechanical biomarkers into clinical studies.

Several limitations should be acknowledged. Only two patient-derived pNF1 cell lines were evaluated; expanding to additional lines and anatomical tumor sites is essential to capture heterogeneity. The 3D hydrogel system does not fully recapitulate the *in vivo* tumor microenvironment, including immune components, vasculature, and native nerve architecture. Future studies integrating organoids, explants, or *in vivo* models will be important. The specific mechanotransduction pathways linking ECM stiffness to intracellular softening, LOX regulation, and drug resistance remain to be elucidated and represent key targets for follow-up investigations.

In conclusion, this work establishes ECM stiffness as a central regulator of pNF1 tumor cell behavior, with direct implications for growth, infiltration, matrix remodeling, and therapeutic response. Understanding stiffness-dependent tumor progression is particularly relevant in the context of surgery, where residual tumors may experience postsurgical ECM stiffening. The study provides a mechanistic framework and a versatile 3D hydrogel platform to investigate pNF1 biology and guide the development of mechano-aware therapeutic strategies, with clear clinical and translational significance.

## Patents

The present study is associated with the U.S. Patent US10227556B2.

## Author Contributions

Conceptualization, K.J.; investigation, K.J. and C. H..; writing-original draft, K.J.; writing-review and editing, K.J. and J. Z.; supervision, K.J. and J.Z. All authors have read and agreed to the published version of the manuscript.

## Acknowledgments

We are deeply grateful to Bonnie F. Sloane, K.J.’s exceptional mentor, who sadly passed away from lung cancer. Her guidance, feedback, and encouragement were instrumental in shaping the NF1 projects and continue to inspire this work. We also sincerely thank K.J.’s NF1 mentoring committee for their ongoing support and mentorship, as well as Raymond Mattingly (East Carolina University Brody Medical School, Greenville, NC, USA), and Harini Sundararaghavan and Katherine Gurdziel (Wayne State University, Detroit, MI, USA) for their valuable input and encouragement through monthly NF1 meetings. Finally, we thank the Neurofibromatosis Therapeutic Acceleration Program (NTAP) for providing funding to support the purchase of immortalized human pNF1 cell lines used in this study.

